# Landscape-scale simulations as a tool in multi-criteria decision making for agri-environment schemes

**DOI:** 10.1101/524181

**Authors:** Chris J. Topping, Lars Dalby, Jose W. Valdez

## Abstract

Increasing concerns over the environmental impacts of agriculture in Europe has led to the introduction of agri-environment schemes (AES) to help mitigate biodiversity loss. However, effectiveness of AES has been mixed and only partially successful in achieving desired outcomes. To improve effectiveness and reduce high costs, multi-criteria decision analysis (MCDA) can help support decision-making and determine the most effective management action. Although MCDA has great potential for evaluating policy measures, it rarely considers the context-dependency of species responses to management practices across different landscapes. Landscape simulations can, therefore, be valuable for reducing the uncertainties when predicting the consequences of management actions. A potential suitable simulation system is the Animal, Landscape, and Man Simulation System (ALMaSS), a mechanistic simulation with can improve MCDA with the automatic integration of a species ecology and behaviour and landscape context. The aim of this study was to demonstrate the effectiveness of ALMaSS in evaluating AES management practices across different landscapes and estimate their ability to achieve the proposed conservation outcomes of three typical species of conservation interest. In this study, the effect of a particular management strategy on a species was dependent on the landscape context, in our case, a combination of landscape structure and the type and distribution of farms, and varied depending on the metrics being measured. Although we did not aim to make recommendations of particular management strategies, we demonstrate how simulations can be used for MCDA to select between management strategies with different costs. Despite the complexity of ALMaSS models, the simulation results provided are easy to interpret. Landscape simulations, such as ALMaSS, can be an important tool in multi-criteria decision making by simulating a wide range of managements and contexts and provide supporting information for filtering management options based on specific conservation goals.

## 1 Introduction

Increasing concerns over the environmental impacts of agriculture in Europe has led to the introduction of agri-environment schemes (AES) which financially compensate farmers for implementing measures that benefit wildlife and help protect the environment (Kleijn and Sutherland, 2003). Some examples of common management actions include crop rotation, introducing buffer strips, managing livestock, rotational set-aside, enhancing habitats for wildlife, and organic farming. Implementation of AES has already led to an increase in the abundance and diversity of a wide range of animal and plant species in areas which may have otherwise not have been afforded such protections (Science for Environment Policy, 2017). However, research suggests that these schemes may only be partially successful in their desired objectives and refinements are necessary to improve effectiveness and reduce their high costs (Kleijn *et al.*, 2001; Berendse *et al.*, 2004; Kleijn *et al.*, 2004; Kleijn *et al.*, 2006; Batary *et al.*, 2015; Żmihorski *et al.*, 2016).

Although the success of any management action will depend upon the goal of management or the policy implemented and the metrics used, there are often multiple objectives, and balancing these with limited budgets can be difficult. Fortunately, AES is adaptive and continually revised, allowing for the refinement of techniques and strategies. One such new approach has been multi-criteria decision analysis (MCDA), a knowledge synthesis methodology which explores potential outcomes of multiple actions according to several criteria (e.g. see Belton and Stewart, 2002; Davies *et al.*, 2013). MCDA’s strength is its ability to handle and analyse the qualitative and quantitative data involved with complex multidimensional environmental decision-making. So far, it has been used to evaluate AES effectiveness in the UK (Park *et al.*, 2004; Westbury *et al.*, 2011), assess long-term environmental impacts of AES on land use in Denmark (Vesterager *et al.*, 2012), and ex post evaluation of AES in the EU (Finn *et al.*, 2009; Vesterager *et al.*, 2012). Although MCDA is a useful tool in providing the foundation for policy decision-making, it has been limited in scope due to the general assumption that the relationship between management action and environmental effects are known. However, this is rarely the case, due to the scarcity of ecological information and the lack of fully recognizing the influence of biotic interactions and spatiotemporal landscape processes.

A major challenge is that even simple landscapes consist of complex ecological networks which make it difficult to accurately predict outcomes and link intervention measures with observed results. A single solution will also rarely encompass all groups of interest and the success of any management scheme may vary between taxa and species (Marshall *et al.*, 2006). For example, although some interventions may benefit a variety of organisms (e.g., species-rich field boundaries can benefit both birds (Vickery *et al.*, 2009) and insect pollinators (Batary *et al.*, 2010)), measures aimed at one taxonomic group or species may be detrimental to others within the ecosystem (e.g., delayed mowing may be beneficial to some bird species, while negatively impacting species which prefer short and sparse ground vegetation (Żmihorski *et al.*, 2016)). Such conflicting results in AES measures can have dire consequences, as illustrated with the local extinction of a butterfly population in the Czech Republic (Konvicka *et al.*, 2008). Consequently, landscape context can also have large and contrasting effects on management outcomes (Tscharntke *et al.*, 2005; Marshall *et al.*, 2006). For example, in landscapes with semi-natural grasslands, local habitat factors were more important than landscape factors in determining the local species richness of butterflies, but in intensively cultivated landscapes, contrary results were found (Ekroos and Kuussaari, 2012). Moreover, in a meta-analysis evaluating the effectiveness of AES, species richness and abundance of pollinator species were found to be primarily driven by landscape context and farmland type, with more positive responses in croplands (vs. grasslands) in simple (vs. cleared or complex) landscapes (Scheper *et al.*, 2013). Understanding how species interact with their environment and the effect of landscape context is thus essential when evaluating policy measures.

Due to such issues, AES have increasingly focused on the landscape scale for improving outcomes, such as providing diverse resources in close proximity through habitat heterogeneity (Benton *et al.*, 2003) and corridors which link habitats for species and enhance biodiversity (Renwick and Lambin, 2011; Delattre *et al.*, 2013). However, determining the precise management action to be undertaken becomes increasingly complex since even small differences in details can produce diverse outcomes. For example, in-field management activities to support AES often involve habitat creation which may vary in shape, type, and management. Although field margin measures will naturally be linear in form, as will within-field overwintering refuge “beetle-banks” (to enhance populations of beneficial arthropods) (Collins *et al.*, 2002), created habitat area is flexible and the width of field margin or plot area can sometimes be a critical factor in determining the number of species it contains. This is partly due to the well-known species-area relationship (e.g. see Ma *et al.*, 2002), but is also a function of the buffering effect against agricultural inputs. In one study, a 3m buffer strip reduced spray drift to the field margin by 95%, while a 6m strip removed it completely (De Snoo and De Wit, 1998). Moreover, the influence of management can also vary depending on the metrics being measured. In the case of grasshoppers, although landscape context determined species diversity, density in field margins was heavily influenced by type of management (Badenhausser and Cordeau, 2012). Making reliable predictions when comparing management actions therefore requires detailed spatial resolution and mechanistic relationships to be accurately represented.

Simulation modelling can be valuable in filling these knowledge gaps and the limitations of MCDA by reducing the uncertainty in predicting the consequences of management actions (Drechsler, 2004). The use of simulation models to evaluate or plan policy implementation has become increasingly more common, ranging from simple models to explain and predict the relationships between policy and effect (e.g. Hailu and Brown, 2007) to complex hierarchical-spatial modelling process approaches (e.g. Rouillard and Moore, 2008). Simulation models have been commonly used in land management issues such as forest planning (e.g. Summers *et al.*, 2015), marine park zoning (Bruce and Eliot, 2006), and to determine ecological and cost-effective solutions in endangered grassland biodiversity (Sturm *et al.*, 2018). However, the wider application of simulation models for policy analysis has been slow due to the conventional use of monitoring and experimental approaches (Parry *et al.*, 2013). For simulation models to be useful as a method to evaluate policy measures, it needs both depth, in terms of detailed mechanistic representation capable of capturing the spatiotemporal interactions common in ecological systems; and breadth, to cover the geographic areas where the policy might be applied.

A potential suitable simulation system for evaluating management policy is the Animal, Landscape, and Man Simulation System (ALMaSS), a mechanistic simulation with can improve MCDA with the automatic inclusion of species modelling and landscape context (Topping *et al.*, 2003). ALMaSS has been already been successfully demonstrated and used in a variety of environmental contexts, including assessment of organic farms on wildlife (Topping, 2011b), energy crop production (Gevers *et al.*, 2011), and recently, to evaluate bidirectional feedbacks between farmer decision-making and land use impacts on wildlife (Malawska & Topping 2017). Although the applicability of simulations, such as ALMaSS models, has been limited by the resource demands of landscape simulation (e.g. hand digitisation of landscapes), this limitation has recently been removed by new methods for handling GIS and data from the EU subsidy schemes, which allows for the rapid creation of highly detailed and accurate agricultural landscapes (Topping *et al.*, 2016). These advancements in geographic data now allows researchers to easily evaluate the influence of landscape-context to management measures for the first time, and open up the potential of landscape-scale simulation models, such as ALMaSS, to be used as a valuable tool in MCDA and policy decision-making.

The aim of this study was to demonstrate the effectiveness of ALMaSS in MCDA by evaluating typical AES management practices in different landscapes and estimate their ability to achieve the proposed conservation outcomes. We determined the benefits of common in-field management strategies for wildlife management of three typical species of interest in agricultural systems in terms of economic return per unit area and assessed the degree to which the return was dependent on landscape-context.

## 2 Methods

### 2.1 ALMaSS

The ALMaSS system is an open source project available on GitLab^1^ and comprises of a set of animal models and an environmental simulator based on different landscapes (Topping, 2017).

#### 2.1.1 Species

The species used in this study represent three typical species considered to be of interest in agricultural systems. The first is the European brown hare (*Lepus europaeus*), a game species which has a national management plan in Denmark due declines since the 1960s (Uldal and Bald, 2013), and is now at low population levels in most of Europe (Strandgaard and Asferg, 1980; Tapper and Parsons, 1984; Smith *et al.*, 2004). The second species is the skylark (*Alauda arvensis*), often considered an indicator species for farmland birds and, due to its decline, the subject of a European management plan (Petersen, 2007). The third species is a common agricultural carabid beetle (*Bembidion lampros*). This species is included because it is widespread and represents an important section of the polyphagous predators carrying out pest control functions in the field (Kromp, 1990; Tamaddoni-Nezhad *et al.*, 2013).

#### 2.1.2 Landscape model

The landscape model provides the environmental context for the animal models. It is a comprehensive environmental simulator with 1-m^2^ resolution and habitats represented by polygons. There are a total of ten different landscapes representing both farmed and non-farmed habitats, with farms being comprised of fields and designated a farm type. This farm type determines the types of crops that are grown, their area coverage, and type of management (see Topping *et al.*, 2016).

#### 2.1.3 In-field management

For the landscape model, we assessed six types of common in-field management:

- Grassy field boundaries (10m wide) with 100%, 50% and 25% of fields applying them (assumed to be cut once in late autumn, October-November)
- Mown field boundaries, in 10m, 5m and 1m widths with 100% of fields applying them (assumed to be cut in May and September)
- Rotational set-aside at 9% by area for pig, arable and mixed rotations
- The addition of field beans to the rotation at 9% by area for pig, arable, and mixed rotations
- Conversion of 25% of conventional farms to organic farms of the same base type
- Unsown patches

The methods for including these in the landscape model differed depending on the type of management. Field boundaries (FB) were created automatically by selecting the fields where these were to be added based on a minimum size (1-ha) and probability (25, 50 or 100%). The boundary was then drawn around the inside of the field removing area from the field. The algorithm for drawing used a pixel-by-pixel stepping method to add rings of 1-m at a time. As a result, complex field shapes sometimes result in failure of the algorithm and failure to add a complete field boundary. Therefore, in the scenario results we use the area covered not the nominal allocation rate for analysis.

Set-aside and field beans were added by including them in the rotation for each farm type such that each farm had 9% of the area under rotational set-aside or field beans. Rotations were assumed to be 110 crops long with the extra 10 (set-aside or field beans) being sensibly placed in the rotation. For field beans, left in the field until the following year, it was important that the following crop was a spring crop with no autumn activities. In this case, spring barley was used to always follow beans.

Unsown patches, also known as skylark scrapes, were added as a management feature to all spring barley and winter-wheat crops grown to maturity (Topping *et al.*, 2013). Thus, whenever a field had these crops present it was also assumed to have skylark scrapes.

### 2.2 Management cost

In order to make decisions regarding the optimal use of management, it is important to know how expensive a management is in terms of economic cost to the farmer, and in terms of practical difficulty. This was achieved by making the assumption that three components were important: the proportion of farm area, the difficulty of the extra machinery cost, and the economic return on the area. The cost index used was calculated as the product of these three metrics (Table 1). For the purpose of this study, the cost indices created is quite arbitrary, because recommendations of particular management strategies is not the aim of this paper. Rather we aim to demonstrate how simulation can be used to select between management strategies with different costs.

**Table 1:**
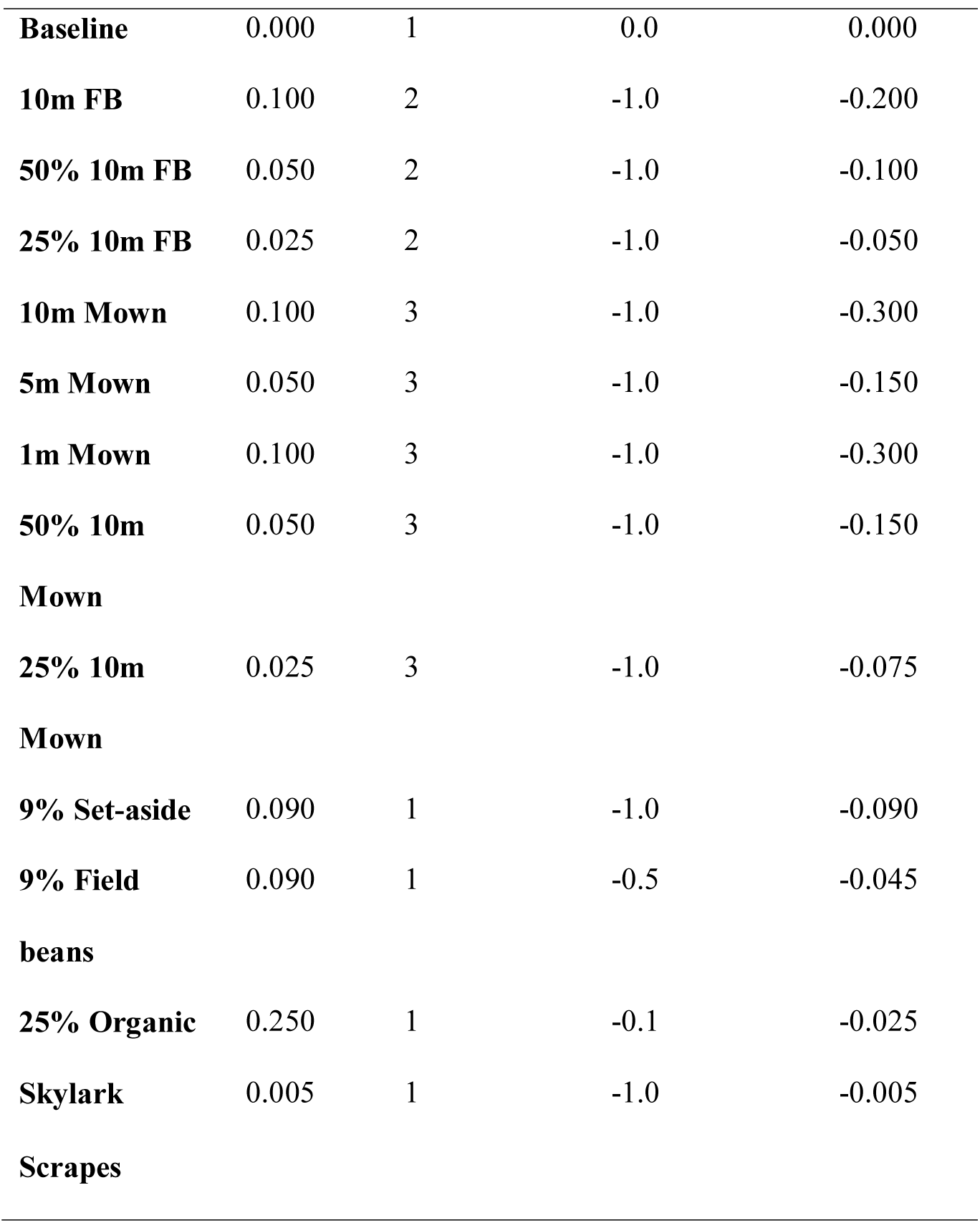
Management strategies and associated cost index assumed for this study based on the product of the area, difficulty and economic effect. Area is in proportion of farmed area; difficulty in extra machinery costs with normal costs being 1.0; and economic effect being an area effect relative to a typical crop.

### 2.3 Simulation scenarios

#### 2.3.1 Animal models

We used three well-established ALMaSS animal models to compare effects using a detailed spatial and temporal resolution (1-m and <1 day), simulating movement, reproduction, and mortality. The European Brown Hare model (Topping *et al.*, 2010) considers hares as individuals represented by detailed foraging and energetics. This model includes detailed foraging and movement behaviour to generate conspecific interactions and activity/intake rates for energetics and breeding based on local habitat quality. The Skylark model (Topping and Odderskaer, 2004; Topping *et al.*, 2013) considers populations of skylarks at an individual level primarily during the breeding season. It includes territory finding, nesting, mating, and foraging behaviour; as well as the energetics of egg and chick development and adult activity. The agricultural carabid beetle model (Bilde and Topping, 2004) considers super-individuals of beetle eggs, larvae, pupae, and female adults. The model includes detailed temperature related development and behaviour in all stages, as well as foraging and hibernation and the associated movement behaviour. In all three models individual animals react to the local presence of conspecifics and also use local information on their environment to make decisions regarding behaviours (e.g. foraging choices, mortality, and dispersal).

All model animals react to direct management conditions. For instance, cutting vegetation will destroy skylark nests and kill infant hares, but only if it is carried out in a place and time where the animals are present. All three models have been used for a number of studies previously (e.g. Topping *et al.*, 2005; Gevers *et al.*, 2011; Topping, 2011b; Parry *et al.*, 2013; Topping *et al.*, 2014; Topping *et al.*, 2015), and detailed documentation about the models may be found in ODdox format (Topping, 2009b, a, 2011a).

#### 2.3.2 Landscapes

Ten landscapes representing a range of Danish conditions and predominant farming types were generated following the methods developed by Topping *et al.* (2016). Farms within the landscape were classified using EU-subsidy information to ensure that farm types and the crops that they grow are up-to-date (although one year old) from The Danish AgriFish Agency (under the Ministry of Food, Agriculture and Fisheries) which maintains an annually updated map of all fields and a database of crops grown in Denmark (“Det Generelle Landbrugsregister” - GLR) (Danish Ministry of Food Agriculture and Fisheries 1999).

Farmers report the crop they intend to grow the following year for each individual field. Data from 2013/14 including contributions from 45,000 farmers was used, making it possible to identify the owner (or manager) of 95% of fields and the actual crop grown on it. During an ALMaSS simulation, crop type on any field at any time is a function of management (crop rotation based on the GLR data), weather, and the soil type of the field. The crop rotation also depends on the farm type, which is created by classifying GLR data together with data on the numbers and type of stock a farm has. This data is available from the Danish Livestock Register (CHR) (Danish Ministry of Food Agriculture and Fisheries 1999), a data set used primarily for disease control in Denmark. A combination of crop and animal information makes it possible to identify major farm types (e.g. pig, arable, or dairy) with less common types, such as sugar beet also identifiable. Additionally, the GLR also indicates whether a farm is organic and its overall size. This extra information provides the basis for a general classification based on logical rules (for details see Topping *et al.*, 2016).

#### 2.3.3 Simulation runs

Each unique combination of the 10 landscapes, 13 management scenarios, and the three species were simulated to create a total of 130 conditions per species. The simulations for each condition were replicated 20 times for each species and mean responses to population changes were calculated in terms of animal density. All scenario runs were for 30 years, with the data analysed based on the mean result of the last 10 years of simulation. To prevent any long-term weather changes influencing the results, weather data were obtained from central Jutland from 1990-1999 and looped three times in the simulations.

#### 2.3.4 Multi-criteria decision making

To implement decision making, clear management goals in terms of both effects and resources required are important. Although there is no single way to determine what management options should be undertaken for wildlife impacts and farm management, two extreme approaches might be to maximise impact or minimise costs. Therefore, for this exercise, we assumed we want to achieve three arbitrary goals whilst minimising costs: i) ensuring we have a minimum of eight adult female hares within km^-2^; ii) avoiding negative impacts on skylark populations of >5%; and iii) increasing beetle numbers by 20%. In order to carry out a management evaluation, the costs of management changes were considered together with the environmental impacts. General costs were calculated for each management using a common metric and used to filter management results.

## 3 Results

### 3.1 Simulation

The baseline densities prior to management varied across landscapes and exhibited different patterns between the three species (Fig. 1). In nearly all cases, the populations of all three species were extant during the last 10 years of simulation. However, long-term declines were evident for hares in Mors, Toftlund and Karup landscapes (Fig. 1). Mean 10-year population densities of all three species were predicted to vary considerably amongst landscapes (Fig. 1).

**Figure 1.**
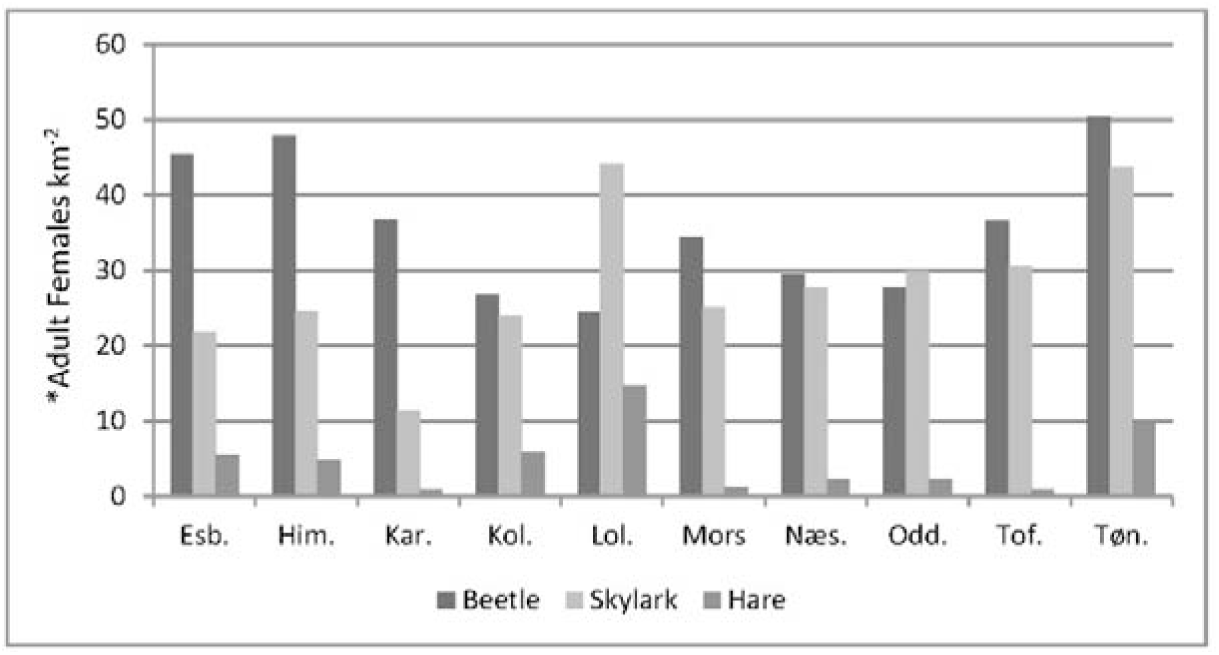
Densities of hares, beetles and skylarks in baseline scenarios for all 10 landscapes considered. *Beetles scaled by 1000. X-axis codes refer to different landscapes: Esb. = Esbjerg, Him. = Himmerland, Kar.=Karup, Kol. = Kolind, Lol.=Lolland, Mors=Mors, Næs.=Næstved, Tof.=Toftlund, Tøn.=Tønder.

The relative impact of any one given management for all three species varies depending on the landscape context (Figure 2). The impact in hares was inversely related to the baseline density for most management, but there was still considerable variation (Figure 2a). For the other species there was no clear density relationship and the pattern of management varies between landscapes and management strategies to a greater extent than the hare (Figure 2). In skylark populations, 10 m mown had negative impacts across all landscapes, especially in Esbjerg, Karup, and Kolind (Fig. 2b). However, skylark scrapes were more positive than other management strategies on skylark (Fig. 2b). In 9 out of 10 cases this management was the most beneficial for skylarks, but in the case of Karup landscape, dominated by potato farmers, the impact was negative (Fig. 2b). The impact of 10m field boundaries and 9% field beans had positive impacts on beetles, especially in half the landscapes with a proportion change of more than 0.3 (Fig. 2b).

**Figure 2.**
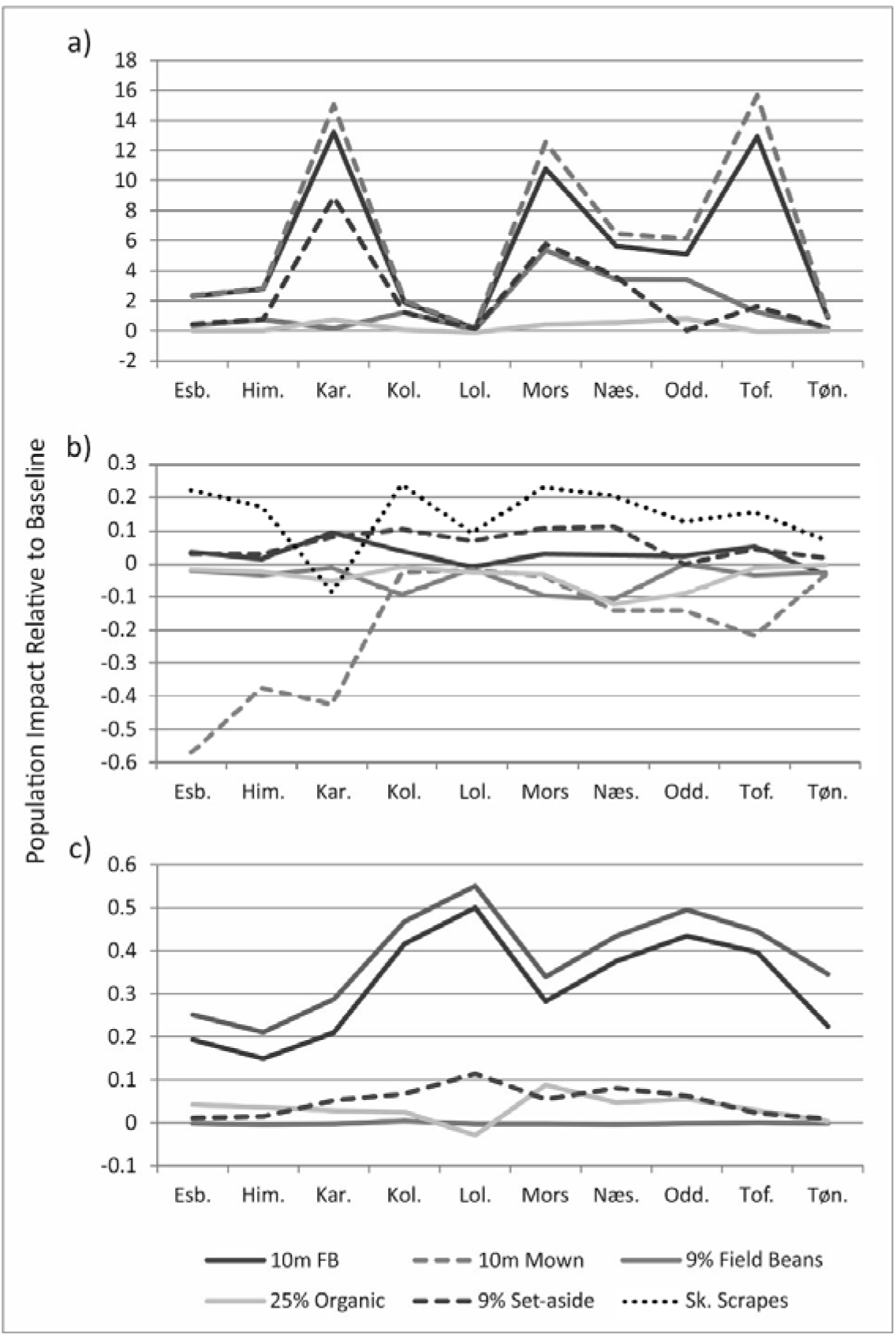
Relative proportion change in population density for different management actions for 10 landscapes. a) Hare; b) Skylark; c) Beetle. X-axis codes refer to different landscapes: Esb. = Esbjerg, Him. = Himmerland, Kar.=Karup, Kol. = Kolind, Lol.=Lolland, Mors=Mors, Næs.=Næstved, Tof.=Toftlund, Tøn.=Tønder.

### 3.2 Multi-criteria decision analysis

The number of possible options where all constraints were met changed as each constraint was added. Using hare constraints only resulted in a mean of 7.9 possible management options per landscape (range 4-13) (Figure 3a), with skylark constraints this was reduced to 5.8 (range 1-12) (Figure 3b), and by adding beetle constraints reduced further still to 2.9 (range 0-7) (Figure 3c). In Himmerland, no management could satisfy all constraints, although beetle increase was only 2% below the target for 10m field boundary management (Figure 3c). Mean costs of achieving the hare management criteria was −0.056 (range 0.00 to 0.09) (Figure 3). Costs approximately doubled by adding beetle and skylark constraints, but the mean cost was still −0.12, lower than the 10m FB management (−0.20), with the range of costs being −0.05 to −0.2 (Figure 3, Table 2). If fewer constraints were applied, the number of management strategies that can be used to satisfy these constraints increases from 29 for all three species to a maximum of 99 for only skylark constraints (Table 2), with the mean costs also decreasing (Figure 3). The 10m field boundaries was the only one that provided consistently good results with all the species combined, but it is expensive in terms of area (Figure 3a,b,c). At the other end of the extreme, doing nothing (zero cost) satisfied only four conditions when hare was a constraint (Figure 3a,b,c,g), but satisfied nearly all the conditions for skylark and skylark with beetle (Figure 3e,f). Compared to the hare only constraint, the beetle only constraint had lower conditions satisfied with only 43 out of 130 (mean 4.3, range 1-7)(Figure 3d) and a cost of −0.025 (Table 2). A skylark only constraint had the most conditions satisfied with 99 out of 130 (mean 4.3, range 6-13)(Figure 3e) and a cost of zero (Table 2), with the baseline producing the best strategy in every single landscape (Figure 3e).

**Table 2.** The number of successful management strategies (N) out of 130 and the mean best cost index across all 10 landscapes for all combinations of species constraints applied to the ten landscapes considered.

| Species constraints | N | Cost Index |
| --- | --- | --- |
| Hare | 79 | -0.056 |
| Hare+Skylark | 58 | -0.066 |
| Hare+Skylark+Beetle | 29 | -0.120 |
| Skylark | 99 | 0.000 |
| Beetle | 43 | -0.025 |
| Skylark+Beetle | 44 | -0.025 |

**Figure 3.**
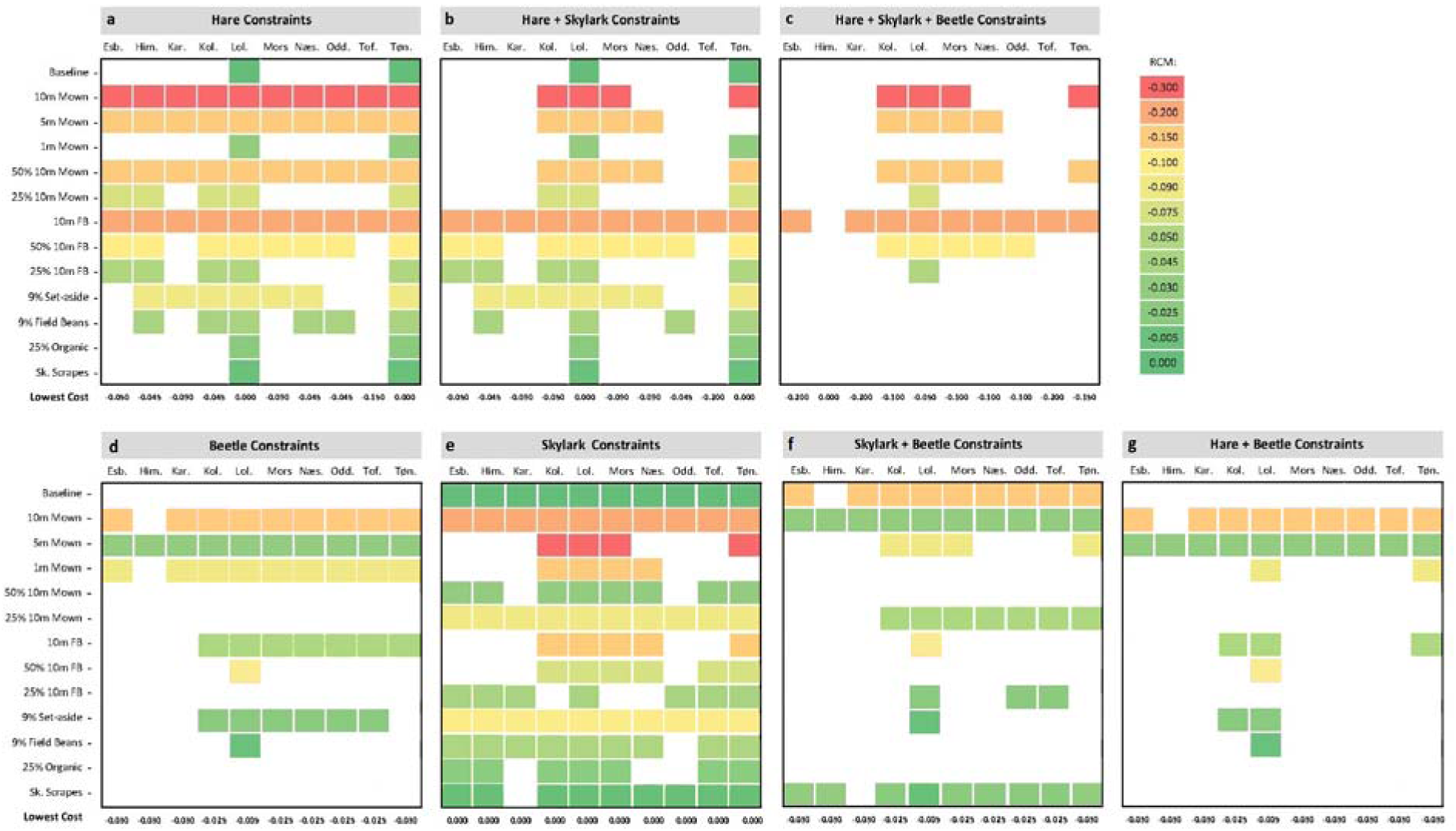
Management options that fulfil the various criteria and their associated costs as an index. White indicates that the criteria were not fulfilled, other colours gradient indicate relative costs of management (RCM). Top codes refer to different landscapes: Esb. = Esbjerg, Him. = Himmerland, Kar.=Karup, Kol. = Kolind, Lol.=Lolland, Mors=Mors, Næs.=Næstved, Tof.=Toftlund, Tøn.=Tønder.

## 4 Discussion

The major outcome of this study is that results were context-dependent in terms of landscape, management, and species responses. All the management strategies tested could be used to achieve a subset of the management goals. Although addition of a 10 metre unmanaged field margin was the most ubiquitously successful management, the impact was highly dependent on landscape for hare and beetle. It is already understood that the precise effects obtained from management measures will be dependent upon the context in which it is applied. For instance, the use of beetle banks to support game-bird populations as a secondary effect of their main function for pest control will only work effectively in areas where game bird resource levels are low, and will not substitute for margins specifically designed for game birds (Thomas *et al.*, 2001). Similarly, provision of habitat or limiting food resources will only benefit rare and declining species in landscapes, where these species still have relict populations (Kleijn *et al.*, 2006). Nearly all management actions will also be more effective in simple agricultural landscapes compared to complex biodiversity-rich landscapes, which will result in little to enhance the ecological processes already present (Tscharntke *et al.*, 2011; Żmihorski *et al.*, 2016). However, in this study, we demonstrate that the effect of a particular management strategy on a species is also dependent on the landscape context, in our case, a combination of landscape structure and the type and distribution of farms. Although the importance of landscape structure and farming was already previously revealed in a meta-analysis on pollinator species (Scheper *et al.*, 2013) and demonstrated from ALMaSS models on hare model densities (Topping *et al.*, 2016), it is clear that this applies to other species as well.

Another context-dependent result was that the responses for each of the three species showed a different pattern to management practices in different landscapes. Considering that the baseline densities for each of the three species differed between landscapes, this result was not a surprise. Similar results have been found on the effects of farm type for agricultural wildlife species, where the benefits of organic farming varied with the landscape structure, and in relation to the species considered (Topping 2011). Similarly, Douglas *et al.* (2009) in their study on the value of field margins as foraging habitats for farmlands, found that management, such as frequent cutting, may be effective in maximizing the benefits for foraging birds by creating short, open vegetation patches. However, frequent cutting does not benefit all groups, with field margins receiving no management or a single July silage cut being able to support greater abundances and species richness of beetles (Woodcock *et al.*, 2007).

This species dependency on management outcomes highlights the need to clearly identify the management desired species-specific goal prior to management implementation. Moreover, whether the results of any management action are deemed successful will depend upon the goal of the management or policy implementing it. In the beetle bank example above, management designed for pest control is likely to fail if assessed as a game bird management. Furthermore, although in-field management prescriptions applied to a relatively small areas can benefit the surrounding landscape (such as greater bumblebee reproduction from sown flower patches (Carvell *et al.*, 2015) or the increase of abundance and species richness in micro- and macro-moth populations from field margins (Fuentes-Montemayor *et al.*, 2011)), this is often not the case, and identifying landscape level affects is required. Thus, two main types of contexts can be identified: the ecological context, pertaining to the effect on biodiversity of a particular management across a landscape, and the policy context pertaining to the goal of the management plan.

Until now, effectiveness of agri-environmental schemes has typically been measured using monitoring and experimental approaches, usually using a general form of index (Braband *et al.*, 2003). Whilst this may describe the general case, it does not account for context-dependency and for the variability of the species responses to different management practices in different landscapes. Moreover, when trying to achieve a specific goal for more than one target species, there will be fewer management strategies capable of producing the target outcome. Considering the context-dependency illustrated by our results and previous studies (Topping 2011; Malawska & Topping 2017), along with the costs associated with AES (Batáry et al. 2015), it is critical to define a clear goal and make efficient use of the available resources when exploring AES management options. During the decision-making process, it is also imperative to decide which management practices will be employed and where, as well as analysing if the costs associated to the chosen management are affordable.

Although in this study we did not aim to make recommendations of particular management strategies, we demonstrate how simulations can be used for MCDA to select between management actions with different costs. Despite the complexity of ALMaSS models, the simulation results provided are easy to interpret and constitute a valuable tool to evaluate in-field management for wildlife across different landscapes. Previously, this would have been of limited usefulness due to the difficulty of generating realistic landscapes and farming scenarios. However, at least for Denmark, this problem has been solved by linking GIS and EU databases to generate landscapes almost automatically. This study demonstrates that ALMaSS, and other similar models, can be an important tool in multi-criteria decision making by simulating a wide range of managements and contexts and provide supporting information to filter management options based on specific conservation goals.

## 5 Conclusions

Although determining the most effective management action is a complex endeavour, an MCDA approach can help support decision-making for policy analysis. However, uncertainties due to the scarcity of ecological information, reduce the confidence to accurately predict the consequences from management actions. This study demonstrates that a species response varies across different landscapes and types of management, and that management outcomes have different affects across landscapes and between species. Landscape simulations models are a valuable tool to help reduce these uncertainties when evaluating wildlife management across different landscapes. ALMaSS and similar mechanistic models can help ensure more cost-effective use of resources, and when considering multi-criteria decision-making, improve location- and goal-specific management outcomes.

## 6 Acknowledgements

We thank Heidi Buur Holbeck, Cammi Aalund Karlslund and Jørn Pagh Berthelsen for invaluable contributions in discussions of the idea behind the current manuscript.

## Footnotes

1 https://gitlab.com/ChrisTopping/ALMaSS_all

